# Engineering Artificial 5′ Regulatory Sequences for Thermostable Protein Expression in the Extremophile *Thermus thermophilus*

**DOI:** 10.1101/2025.05.06.652417

**Authors:** Che Fai Alex Wong, Shizhe Zhang, Lisa Tietze, Gurvinder Singh Dahiya, Rahmi Lale

**Affiliations:** Department of Biotechnology, Faculty of Natural Sciences, Norwegian University of Science and Technology, Trondheim, Norway; Syngens AS, Trondheim, Norway

**Keywords:** Extremophiles, *Thermus thermophilus*, Escherichia coli, artificial promoter, thermostable protein expression

## Abstract

The utilisation of biocatalysts in biotechnological applications often necessitates their heterologous expression in suitable host organisms. However, the range of standardised microbial hosts for recombinant protein production remains limited, with most being mesophilic and suboptimal for certain protein types. Although the thermophilic bacterium *Thermus thermophilus* has long been established as a valuable extremophile host, thanks to its high-temperature tolerance, robust growth, and extensively characterised proteome, its genetic toolkit has predominantly depended on a limited set of native promoters. To overcome this bottleneck, we have expanded the available regulatory repertoire in *T. thermophilus* by developing novel artificial 5′ regulatory sequences.

In this study, we applied our Gene Expression Engineering platform to engineer 53 artificial 5′ regulatory sequences (ARES) in *T. thermophilus*. These ARES, which comprise both promoter and 5′ untranslated regions (UTRs), were functionally characterised in both *T. thermophilus* and *Escherichia coli*, revealing distinct host-specific expression patterns. Furthermore, we demonstrated the utility of these ARES by demonstrating high-level expression of thermostable proteins, including *β*-galactosidase, a superfolder citrine fluorescent protein, and phytoene synthase. A bioinformatic analysis of the novel sequences has also being carried out indicating that the ARES possess markedly lower GC content compared to native promoters. This study contributes to expanding the genetic toolkit for recombinant protein production by providing a set of functionally validated ARES, enhancing the versatility of *T. thermophilus* as a synthetic biology chassis for thermostable protein expression.

## Introduction

Evolution has generated a vast array of biocatalysts across diverse biological systems. Recent advances in metagenomics have begun to reveal this largely untapped diversity, opening new avenues for biotechnological applications^1^. However, to harness these biocatalysts it is often necessary to express them heterologously in host organisms that can produce them in sufficient quantities while maintaining their functional integrity^2^. Although several microbial hosts have been standardised for industrial and environmental applications^3^, the repertoire is predominantly limited to mesophilic hosts. This limitation is particularly pronounced when expressing proteins that require the robust folding and stability provided by high-temperature environments^4^.

The thermophilic bacterium *Thermus thermophilus* is a unique extremophile. This Gram-negative organism, which has a genomic GC content of approximately 69%^5^, thrives at temperatures between 55 °C and 80 °C. Its thermostable enzymes such as DNA polymerase are already widely used in biotechnological processes^6^. In addition, nearly 20% of its proteins have been structurally characterised by X-ray crystallography, making it one of the most well-studied thermophiles. Key advantages include high cell densities in simple media, an aerobic metabolism that simplifies laboratory handling, and relatively high natural transformation efficiencies^7^.

The number of well characterised promoters for *T. thermophilus* has been limited. To overcome this bottleneck, we employed our Gene Expression Engineering (GeneEE) platform to create artificial 5′ regulatory sequences (ARES) that do not rely on prior knowledge of native promoter architecture^8^. In our previous work, we demonstrated the feasibility of the GeneEE approach in seven different microorganisms^8,9^. In this study, we report the establishment of 53 ARES in *T. thermophilus* and characterisation in both *T. thermophilus* and *E. coli*. We further validate these regulatory elements by driving the high-level expression of thermostable enzymes, thereby underscoring the potential of these synthetic parts for enhanced expression relevant for biotechnological applications.

## Materials and Methods

### Bacterial Strains and Growth Conditions

*E. coli* DH5*α* was used for both cloning and expression. Cultures were grown in Lysogeny Broth (LB; tryptone 10 g/L, yeast extract 5 g/L, NaCl 5 g/L) supplemented with 50 mg/L kanamycin when necessary. For the screening of ARES and expression studies, a knockout strain of *Thermus thermophilus* HB27 with high natural transformation efficiency (HB27Δago, genome accession number: AE017221) was used. *T. thermophilus* cultures were grown in Thermus Broth (TB; bactotryptone 8 g/L, yeast extract 4 g/L, NaCl 3 g/L), prepared in commercial mineral water (Farris, Ringnes AS, Oslo), and supplemented with 30 mg/L kanamycin when selection was required. Overnight cultures of *E. coli* were incubated at 37 °C, whereas *T. thermophilus* cultures were grown at 65 °C with agitation at 150 rpm. For solid medium cultivation, TB agar plates were incubated in a sealed, damp plastic box to prevent drying. Due to occasional colony merging on solid TB, only isolated colonies were selected for further experiments. For storage, an overnight culture of *T. thermophilus* was centrifuged at 5000 xg for 5 min at room temperature, the supernatant removed, and the cell pellet stored at –20 °C. Prior to experiments, pellets were thawed at room temperature, resuspended in 1 mL of TB, and then inoculated into 20 mL of pre-warmed TB in 100–150 mL baffled Erlenmeyer flasks.

For transformation into *T. thermophilus*, we followed the protocol provided by Dr. José Berenguer’s group. Briefly, an overnight culture was diluted 1:50 in fresh pre-warmed TB and incubated at 65 °C until reaching log phase (OD_550_ ~0.4). Then, 0.8 mL of culture was transferred to a 12 mL tube and mixed with approximately 300 ng of plasmid DNA. After a further 4-hour incubation at 65 °C with shaking, the cells were plated on pre-warmed selective TB agar. Colonies typically appeared overnight; however, plates were maintained for additional nights to allow slower-growing transformants to emerge.

### Plasmid DNA Library Construction

A shuttle plasmid, pMK184^10^, was used for constructing the DNA libraries. The plasmid comprises four parts: (i) a dual replicon backbone; (ii) a kanamycin resistance marker; (iii) a reporter gene encoding a superfolder citrine fluorescent protein (IFP) from pMoTK110^11^; and (iv) a GeneEE segment that is 200 nt long with random nucleotide composition^8^. These segments are flanked by primer binding sites (BioBrick Prefix and Suffix, see Supplementary Table S1 for primer sequences) which enable conversion from single-stranded oligonucleotides (synthesised by Integrated DNA Technologies, Coralville, Iowa, USA) to double-stranded DNA and facilitate cloning via Gibson or Golden Gate assembly (Figure 1A).

**Figure 1.**
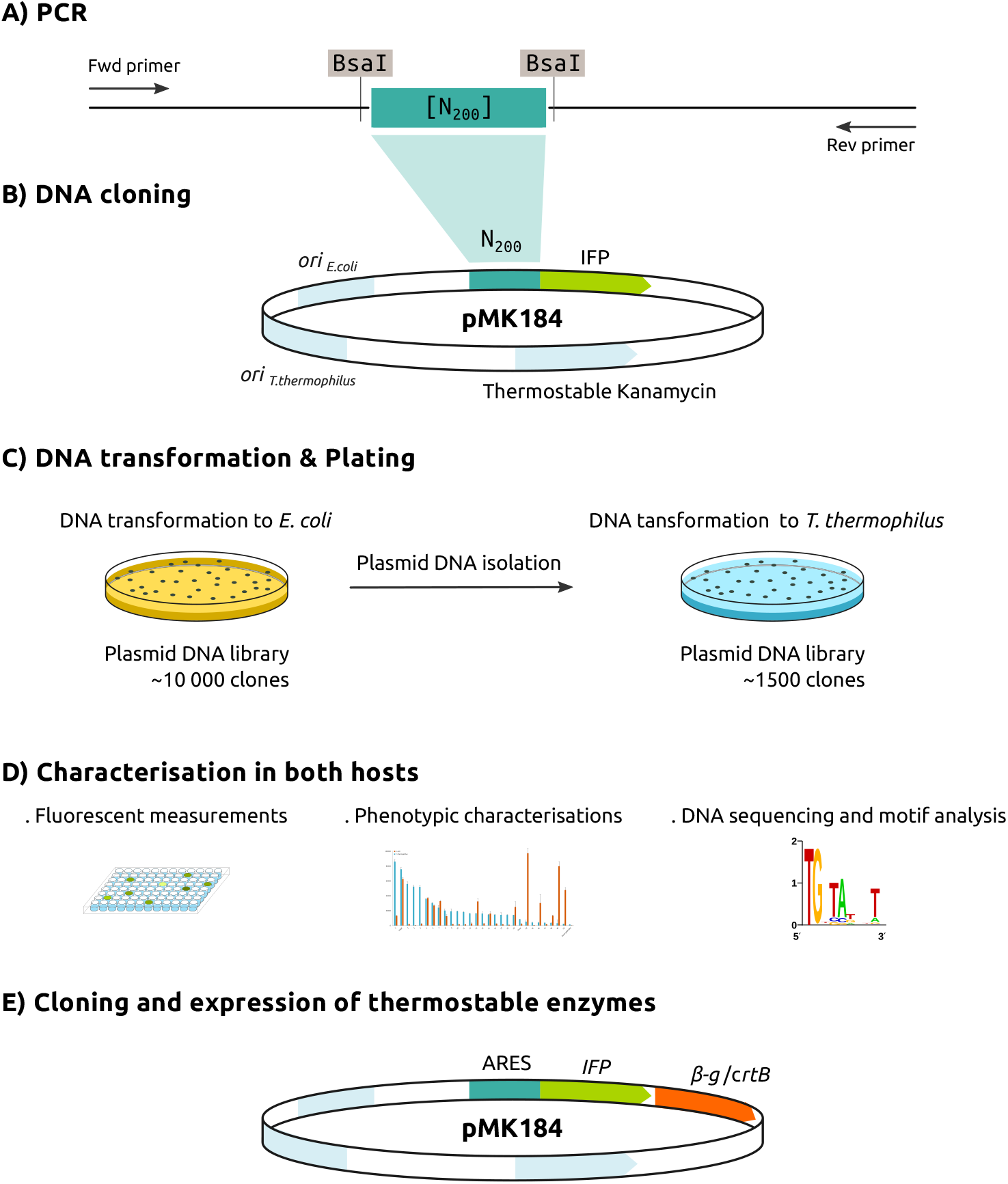
Schematic overview of the study. The plasmid library was constructed by cloning a 200-nucleotide random sequence upstream of the thermostable reporter gene, superfolder citrine fluorescent protein (IFP). Plasmid DNA library was pooled from *E. coli* and transformed to *T. thermophilus*. Approximately 1,500 *T. thermophilus* clones, arrayed on 96-well plates, were screened under UV light, and 53 fluorescent clones were selected for further characterisation. The thermostable enzymes were transcriptionally fused to the IFP CDS. Plasmid DNAs were subsequently transformed into *E. coli* for comparative expression analysis.

Two plasmids were constructed as positive controls: one with the strong promoter *P*_*slpA*_, and one with the medium-strength promoter *P*_*nqo*_, both driving IFP expression^11^ (plasmid maps are available both in the Supplementary and in our [GitHub repository](https://github.com/LaleLab/ares_thermus)). The backbone pMK184 was linearised using back-to-back PCR with primers Tt1 and Tt2. *P*_*slpA*_ was amplified from pMK184 using primers Tt3 and Tt4, and IFP was amplified from pMoTK110 with primers Tt5 and Tt6. After DpnI digestion to remove parental plasmid and purification (QIAGEN PCR purification kit), the fragments were assembled using Gibson cloning^12^. The resulting plasmid (pMK184-Pslp-IFP) was verified by DNA sequencing. To facilitate subsequent Golden Gate cloning, the BsaI sites in pMK184-Pslp-IFP were removed via overlapping PCR (primers Tt7–Tt10) and reassembled by Gibson cloning.

The second control plasmid, pMK184-Pnqo-IFP, was constructed from the BsaI-free pMK184-Pslp-IFP by replacing the *P*_*slpA*_ promoter with *P*_*nqo*_ (amplified using primers Tt11 and Tt12, with exclusion of *P*_*slpA*_ via primers Tt13 and Tt14).

For ARES library construction, pMK184-Pslp-IFP was PCR amplified with primers Tt15 and Tt16 to excise the *P*_*slpA*_ region and introduce BsaI sites. After DpnI digestion and purification, the randomised GeneEE segments (75 ng total) were ligated into the backbone via Golden Gate assembly (30 cycles of alternating 37 °C and 16 °C for 5 min each, final inactivation at 65 °C for 20 min)^13^ (Figure 1B). The resulting library was transformed into *E. coli* DH5*α*, yielding approximately 10,000 colonies. Plasmids were then isolated and used for transformation into *T. thermophilus*; however, due to colony merging on TB agar, only 1,500 individual *T. thermophilus* colonies could be recovered (Figure 1C).

### Selection and Characterisation of ARES Strength

Using IFP as a reporter, an initial visual screening under UV light was performed on the 1,500 *T. thermophilus* clones (Figure 1D). Fifty-three clones displaying fluorescence were selected (yielding a 3.5% hit rate) and cultured overnight in 1.5 mL TB in 96 deep-well plates at 65 °C. The next day, a 96-pin replicator was used to transfer cultures into separate plates for storage at –20 °C and for plasmid isolation.

For quantitative assessment, cells were pelleted (three biological replicates) and resuspended in an equal volume of phosphate-buffered saline (PBS). One hundred microlitres of each suspension was transferred to 96-well black microtitre plates. Fluorescence (excitation: 485 nm, emission: 520 nm) and OD_600_ were measured using a TECAN SpectraMax reader, and fluorescence values were normalised to cell density. Plasmids from the 53 clones were then isolated (QIAGEN protocol) and Sanger sequenced using primer Tt17 to determine their ARES sequences (See Supplementary Table S2).

The same 53 plasmids were transformed into *E. coli* for cross-characterisation. Cultures were grown overnight in 1.5 mL LB at 37 °C, washed and resuspended in PBS, and then analysed for normalised fluorescence.

### Cloning and Expression of Thermostable *β*-Galactosidase and Phytoene Synthase Using ARES

To evaluate the functionality of the isolated ARES for recombinant protein production, two thermostable enzymes—*β*-galactosidase and phytoene synthase—were selected as model proteins^11^. The gene variant encoding *β*-galactosidase (denoted as *β-g*) and the gene for phytoene synthase (*crtB*) (see Supplementary Table S3) were cloned by transcriptionally fusing to the IFP gene (Figure 1E) as follows. Initially, the *β*-g gene was amplified to include BamHI and PvuI restriction sites using primers Tt18 and Tt19, while the *crtB* gene was amplified with HindIII and PvuI sites using primers Tt20 and Tt21. A pooled plasmid preparation, consisting of all 53 plasmids each harbouring a unique ARES, was then digested with either BamHI-HF and PvuI-HF or HindIII-HF and PvuI-HF to establish the corresponding ARES libraries. The digested *β-g* and *crtB* fragments were ligated into the appropriate ARES library backbones and transformed into *E. coli*. After scraping all *E. coli* transformants from the agar plates, the plasmid DNA was isolated and subsequently transformed into *T. thermophilus*. For each gene, 12 transformants exhibiting fluorescence under UV illumination were selected for further experiments.

### Measurement of *β*-Galactosidase Activity

Cell-free extracts were prepared using CelLytic B (Sigma-Aldrich) with 0.2 mg/mL lysozyme. Briefly, 1.5 mL of *T. thermophilus* culture (OD_550_ ~0.6) was centrifuged at maximum speed for 2 min. The pellet was resuspended in 400 *µ*L of 2X CelLytic B solution and vortexed for 10 min. After centrifugation (5 min at max speed), the supernatant was used for enzyme assays.

*β*-Galactosidase activity was measured using *o*-nitrophenyl-*β*-*D*-galactopyranoside (ONPG) as a substrate. A 5 mM ONPG solution in 50 mM sodium phosphate buffer (pH 7.0) was prepared, and 20 *µ*L of cell extract was mixed with 80 *µ*L ONPG in a transparent 96-well plate. After incubation at 65 °C for 15 min, the reaction was terminated by cooling on ice for 5 min and adding 100 *µ*L of 0.5 M Na_2_CO_3_. The reaction product *o*-nitrophenol (ONP) was quantified by measuring absorbance at 420 nm, with protein concentration normalised using absorbance at 280 nm.

### Extraction and Absorbance Measurements of Carotenoids

For phytoene synthase expression, 3 mL of *T. thermophilus* culture (OD_550_ ~3.0) was centrifuged at full speed for 5 min. The pellet was resuspended in 1 mL acetone and sonicated for 5 min, followed by centrifugation (5 min at full speed). An absorbance scan from 400 to 500 nm was performed using 100 *µ*L of the acetone extract in an Infinite M200 Pro TECAN fluorimeter. The absorbance at 452 nm was recorded as an indicator of carotenoid production^14^.

### IFP Fluorescence Measurements of Clones with Enzyme Expression

IFP fluorescence was measured in *T. thermophilus* transformants carrying plasmids for enzyme expression. Individual colonies were grown overnight in 4 mL LB. The next day, 1 mL of culture was centrifuged at maximum speed, and the pellets were resuspended in an equal volume of PBS. Both fluorescence (excitation: 485 nm, emission: 520 nm) and OD_600_ were measured, and the values were normalised to assess expression levels.

### Statistical Analysis

One-tailed Student’s t-tests were performed to compare fluorescence values and enzymatic activities among different ARES. Significance is indicated in the figure legends with asterisks denoting p-values: *** (p < 0.001) and ** (p < 0.05) when compared to the positive control, and +++ or ++ when compared to the negative control.

For correlation analysis, ARES were ranked by fluorescence intensity in both *T. thermophilus* and *E. coli* (rank 1 for highest expression to 53 for lowest). The linear correlation coefficient between these rankings was calculated, and the corresponding p-value is provided in the relevant figure legend.

### Genomic Data Analysis

Genomic datasets for *Escherichia coli* DH5*α* (accession number CP080399) and *Thermus thermophilus* HB27 (accession number GCA_000008125) were downloaded from EMBL. GenBank files were parsed using BioPython to extract coding sequences (CDS) and the upstream intergenic regions (IRs). These sequences were stored in parquet format and made available in our GitHub repository. GC contents were calculated for both CDSs and IRs.

Additionally, RNA-seq datasets (for *E. coli* DH5*α*: Run Accession SRR10995619-21, and for *T. thermophilus*: Run Accession SRR1038512-17) were processed using Salmon^15^ to obtain normalised Transcripts Per Million (TPM) values. CDS, IR sequences, and TPM values were merged into a dataframe, and genes were categorised into high (>90% TPM), medium (50–90% TPM), and low (<50% TPM) expression buckets for further analysis.

## Results

### Isolation of 53 Artificial Regulatory Sequences in *T. thermophilus*

To generate artificial 5′ regulatory sequences in *T. thermophilus*, we replaced the native promoter of the reporter gene, superfolder citrine fluorescent protein (IFP), with a 200-nucleotide GeneEE fragment (Figure 1A, B) in the shuttle plasmid, pMK184^10^. The initial ARES plasmid DNA library, generated in *E. coli*, comprised approximately 10,000 clones. After transformation of the library into *T. thermophilus*, 1,500 individual colonies were obtained. Visual screening under UV light identified 53 fluorescent clones (3.5% hit rate), which were then cultured in 96-deep-well plates for further analysis (Figure 1C, D).

To benchmark expression strength, two native *T. thermophilus* promoters were used as controls: (i) *P*_*slpA*_, a strong cross-species promoter; and (ii) *P*_*nqo*_, a medium-strength promoter with reduced activity in *E. coli*. In *T. thermophilus*, the ARES clones exhibited a broad range of fluorescence intensities (Figure 2). The strongest ARES clone 1 exceeded the fluorescence of the strong *P*_*slpA*_ control, while the majority (49/53) showed expression levels within ±1-fold of *P*_*nqo*_. Notably, ARES clones 1–4 displayed minimal background expression in *E. coli*, which is advantageous when cloning potentially growth inhibiting proteins in *E. coli*. When comparing the normalised

**Figure 2.**
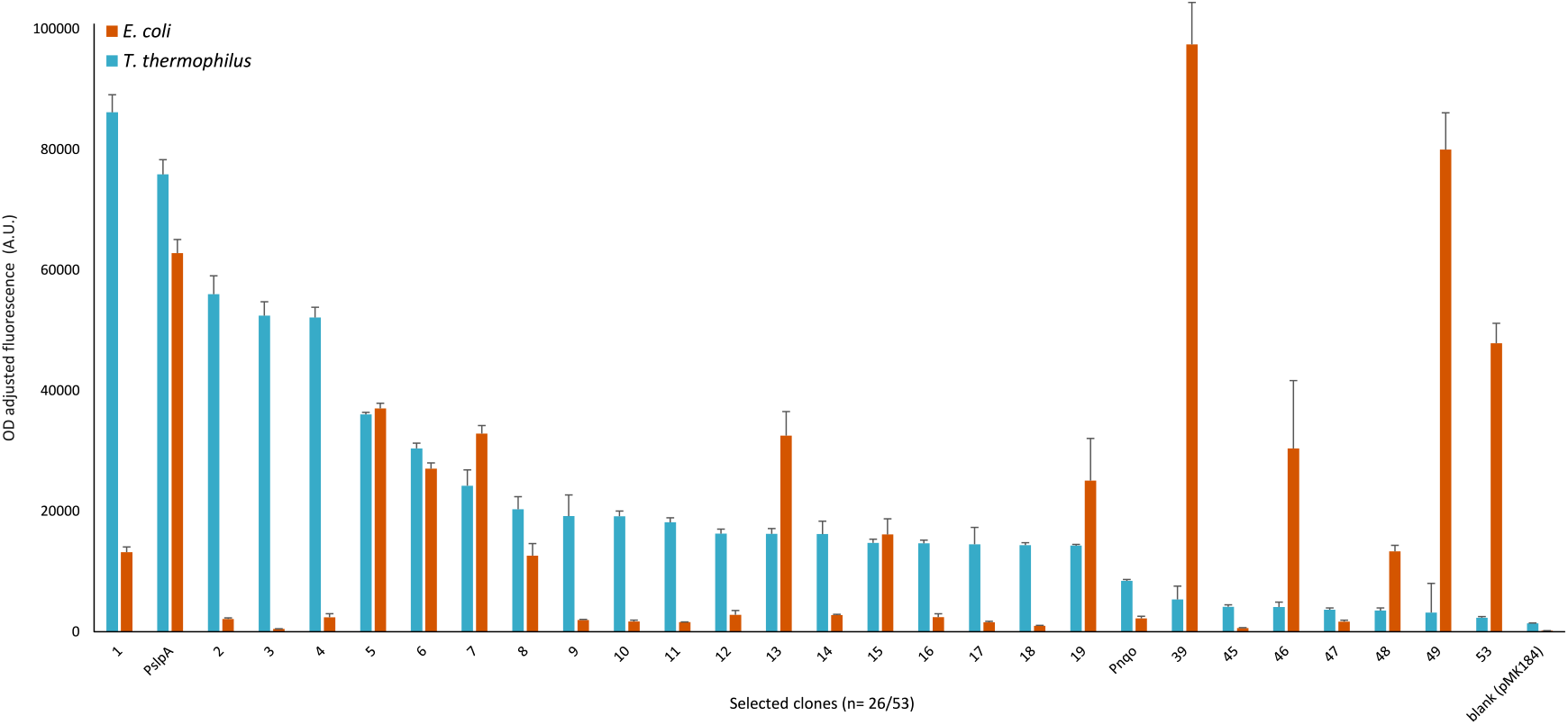
Normalised IFP fluorescence of the isolated ARES in *T. thermophilus* (orange) and *E. coli* (blue). (a) The isolated ARES exhibit a range of expression strengths, with the strongest clone (ARES 1) exceeding the fluorescence of the strong control *P*_*slpA*_, in *T. thermophilus*. (b) Most ARES (49/53) display expression levels similar to the medium control *P*_*nqo*_. (c) In *E. coli*, a similar trend is observed; however, clones 1–4 show notably lower expression, which is advantageous for cloning toxic genes. Only 25 representative clones are shown in the figure; for all clones and their phenotypes see Supplementary Table S2. Positive controls: *P*_*slpA*_, and *P*_*nqo*_. Negative control: empty plasmid pMK184. fluorescence intensity values between the two hosts, 45 of 53 clones exhibited stronger expression in *T. thermophilus* than in *E. coli*, underscoring the benefits of performing a functional screening directly in the target (thermophilic) host.

### DNA Sequence Analysis

Sanger sequencing was performed on all 53 ARES clones (see Supplementary Table S2 for the DNA sequences). An analysis of the GC content revealed that, despite the high genomic GC content of *T. thermophilus* (approximately 69%, Table 1), regulatory regions are typically AT-enriched likely to facilitate DNA strand separation during transcription. In our study, while the native promoters *P*_*nqo*_ and *P*_*slpA*_ have GC contents of approximately 60%, the ARES had an average GC content of 44% (±6%), ranging from 29% (ARES 52) to 73% (ARES 41). In line with other GeneEE studies, these results underscore the flexibility of the bacterial transcriptional machinery in accommodating a wide variety of regulatory sequences. Moreover, none of the ARES did yield significant hits in BLAST searches (no significant hits found using tblastn as of April 2025), suggesting that they represent novel sequence solutions that are rarely or if at all encountered in this bacterium.

**Table 1.** Comparative genomic features of *E. coli* and *T. thermophilus*. Gene density (%) represents the percentage of the genome occupied by annotated genes, whereas IR density (%) denotes the fraction occupied by intergenic regions. GC content is presented for the entire genome, genes, and IRs.

|  | *E. coli* | *T. thermophilus* |
| --- | --- | --- |
| Genome size | 4,490,755 | 1,894,877 |
| Number of genes | 4,205 | 1,969 |
| Sum of genes length | 3,961,399 | 1,796,235 |
| Sum of IRs* | 529,356 | 98,642 |
| Gene density (%) | 88.21 | 94.79 |
| IR density (%) | 11.79 | 5.21 |
| GC content (%) |  |  |
| Genome | 50.74 | 69.44 |
| Genes | 51.80 | 69.59 |
| IRs | 47.35 | 68.76 |
| IFP |  | 41.94 |

\* IR, intergenic region.

### Characterisation of Enzyme Expression Using ARES

To validate the functionality of the ARES for driving expression of biotechnologicaly relevant thermostable enzymes, we transcriptionally fused thermostable *β*-galactosidase and phytoene synthase by placing them downstream of the IFP CDS (Figure 1E). For each enzyme, 12 fluorescent clones (each under the control of a different ARES; see Supplementary Table S2) were selected.

Initially, IFP fluorescence intensities were measured on enzyme-expressing transformants. Clones expressing *β*-galactosidase or phytoene synthase under different ARES showed fluorescence profiles correlating with the enzymatic activities measured (Figures 3 and 4). This correlation supports the utility of the ARES in recombinant gene expression with transcriptionally-fused thermostable proteins.

**Figure 3.**
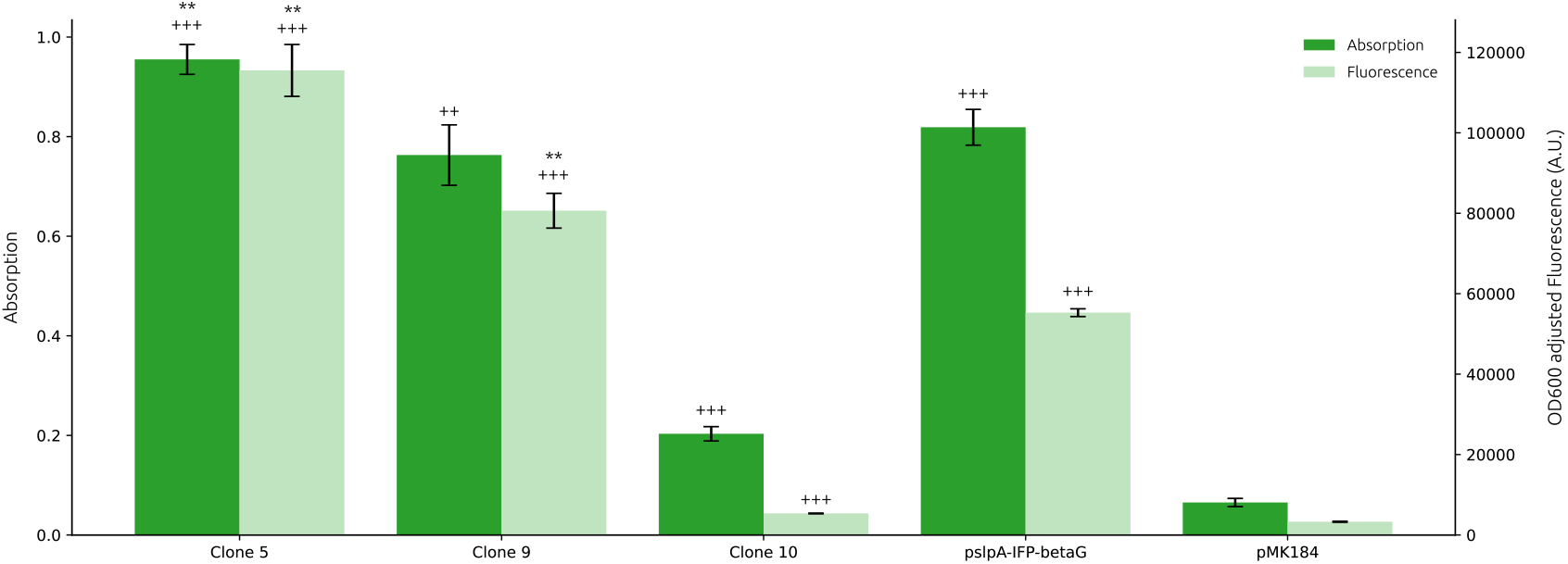
Absorbance and fluorescence measurements of *β*-galactosidase and IFP expression in *T. thermophilus* clones. Clones 5, 9, and 10 displayed *β*-galactosidase activity using the substrate *o*-nitrophenyl-*β*-*D*-galactopyranoside. These clones represent *T. thermophilus* colonies transformed with ligated plasmids carrying three different ARES for driving *β*-galactosidase expression. The positive control (pslpA-IFP-betaG) is a transformant harbouring a plasmid with the strong promoter *P*_*slpA*_, while the negative control (pMK184) contains the native plasmid backbone without an inserted promoter. Absorbance values represent mean ± s.d. of n = 3. *** indicates a p-value < 0.001 compared to the positive control; ** indicates a p-value < 0.05 compared to the positive control; +++ indicates a p-value < 0.001 compared to the negative control; ++ indicates a p-value < 0.05 compared to the negative control. Fluorescence measurements of IFP expression in the same clones showed a similar pattern. Clone 5 exhibited higher IFP fluorescence than the positive control, indicating elevated expression; clone 9 also demonstrated higher expression, while clone 10 showed reduced IFP fluorescence. Fluorescence values represent mean ± s.d. of n = 3. *** indicates a p-value < 0.001 compared to the positive control; +++ indicates a p-value < 0.001 compared to the negative control.

**Figure 4.**
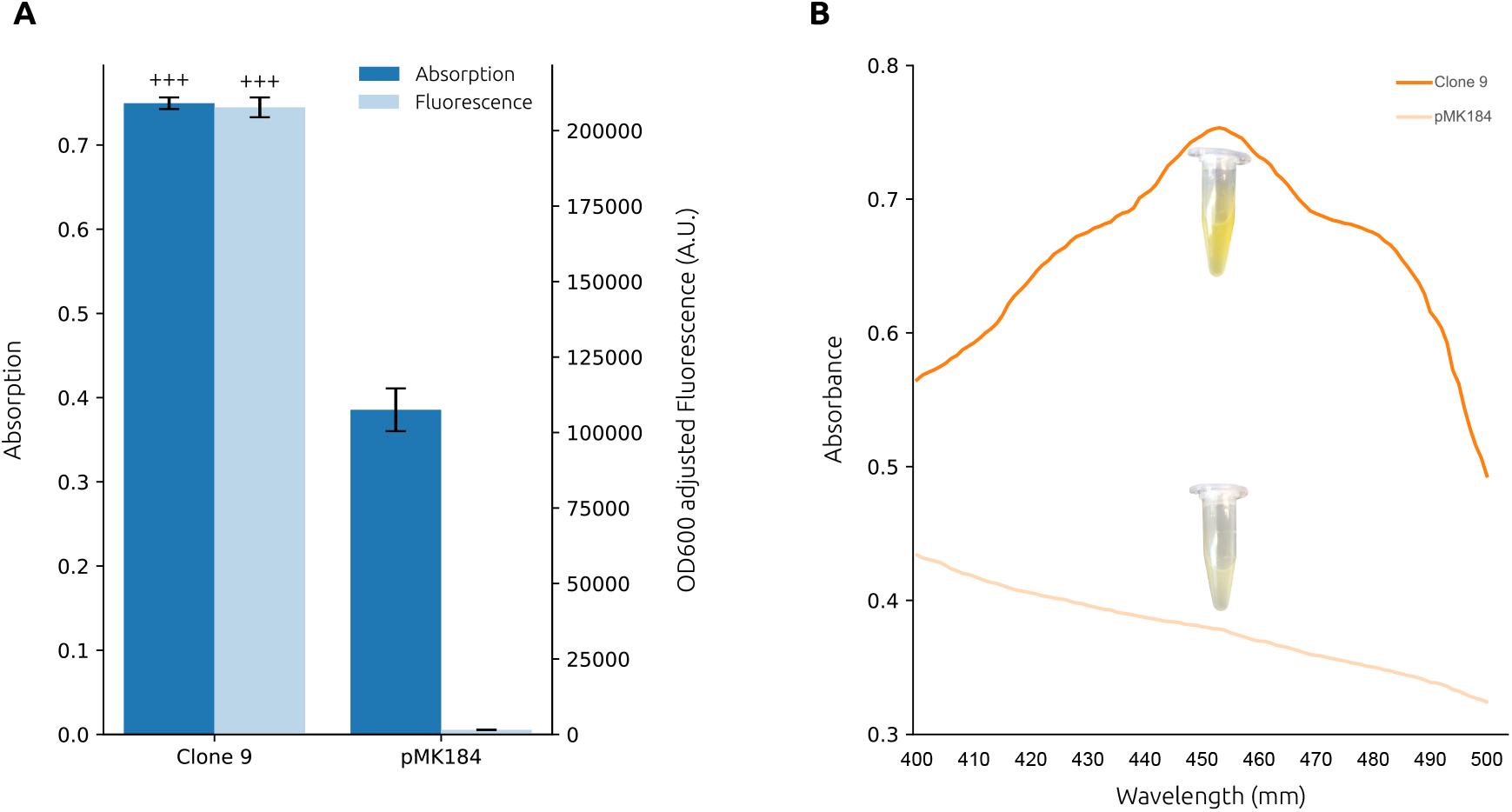
Absorbance and fluorescence assessments of phytoene synthase and IFP expression in *T. thermophilus* clone. Clone 9 represents *T. thermophilus* colony transformed with a ligated plasmid carrying an ARES driving both IFP and phytoene synthase expressions, while pMK184 as the negative control represents the transformant carrying the empty plasmid backbone. The expression of phytoene synthase is represented by the presence of carotenoids from absorbance measurements (A and B) and visual evaluation (B). Bar graphs (A) display absorbance measured at 452 nm and fluorescence values for Clone 9 and pMK184. Clone 9 shows higher absorbance and fluorescence compared to pMK184, indicating the expression of both phytoene synthase and IFP. Absorbance and fluorescence values represent mean ± s.d. of n = 3. +++ indicates a p-value < 0.001 compared to the negative control. The orange colour of the culture and the distinct absorption peak at 452–453 nm (B) further indicate carotenoids production in clone 9.

*β*-Galactosidase activity was assessed via *o*-nitrophenyl-*β*-*D*-galactopyranoside (ONPG) conversion. Three clones exhibited detectable enzymatic activity: clone 5 showed significantly higher activity than the positive control (*P*_*slpA*_), clone 9 was comparable to the positive control, and clone 10 had lower activity (Figure 3). These differences confirm that the ARES can be used to obtain a range of expression levels even when fused to a fluorescent protein.

For phytoene synthase, enzyme function was evaluated both visually and spectrophotometrically (Figure 4). The culture of clone 9 displayed an orange colour, in contrast to the pale yellow of the negative control (pMK184) (Figure 4B). An acetone extract of clone 9 showed an absorbance peak at 452–453 nm (Figure 4B), consistent with carotenoid production and indicative of a *β, β*-carotene type chromophore^14,16^. These findings demonstrate that the isolated ARES are suitable for engineering both reporter and functional enzyme expression in *T. thermophilus*.

## Bioinformatic Analysis

To contextualise the performance of ARES in *T. thermophilus* versus *E. coli*, we conducted a comparative bioinformatic analysis of both genomes, focusing on overall genomic features, gene length, intergenic region (IR) length, and GC content. Table 1 summarises key statistics, including genome size, number of genes, total gene length, IR length, and average GC content for each organism.

As shown in Table 1, *E. coli* has a larger genome (4.49 Mb) with more genes (4,205) than *T. thermophilus* (1.89 Mb, 1,969 genes). Correspondingly, *E. coli* exhibits lower gene density (88.21%) and higher IR density (11.79%) compared to *T. thermophilus* (94.79% and 5.21%, respectively), indicating a more compact genome in the thermophile. Notably, the average GC content is significantly higher in *T. thermophilus* (69.44%) than in *E. coli* (50.74%), a likely consequence of adaptations to high-temperature environments.

Figure 5 shows the distribution of gene lengths in *T. thermophilus* (blue) and *E. coli* (orange). Both species cover a similar spectrum of gene sizes, with gene length extending beyond 5,000 bp, indicating that thermophilic genome streamlining has not substantially altered gene length distributions. Notably, among the top 10 % most highly expressed genes (per our RNA-seq analysis; see Materials and Methods), *T. thermophilus* genes tend to be marginally shorter compared to their counterparts in *E. coli* (Figure 5B).

**Figure 5.**
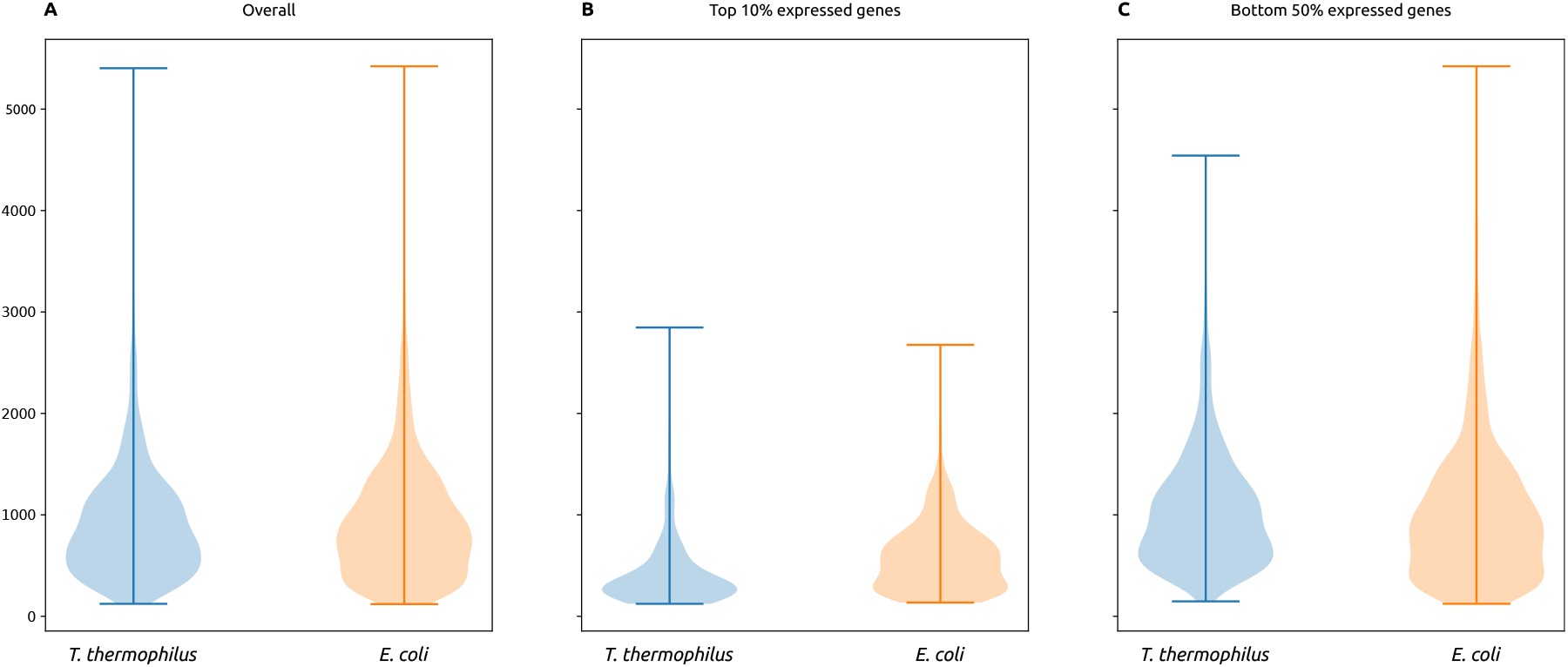
Genelength distributions in *T. thermophilus* (blue) and *E. coli* (orange). Violin plots show overall gene lengths (A), lengths for the top 10% of highly expressed genes (B), and lengths for the bottom 50% of weakly expressed genes (C). The y-axis indicates gene length in base pairs.

Figure 6 shows the GC content distributions for each organism’s overall gene set at the genome level (Figure 6A), as well as for the top 10% (Figure 6B) and bottom 50% of expressed genes (Figure 6C). *T. thermophilus* (green) retains its high-GC signature across all categories, whereas *E. coli* (red) clusters around 50–51%. Notably, the synthetic ARES (blue and orange) have lower GC content than the native genomic regions, consistent with the observation that many functional promoters and regulatory elements are AT-enriched, potentially aiding transcriptional initiation. Finally, Figure 7 depicts the distributions of intergenic region (IR) lengths in *T. thermophilus, E. coli*, and among the ARES constructs. At the genome level, *T. thermophilus* (green) displays a much broader IR length range, exceeding 3,500 bp in some cases, whereas *E. coli* (red) remains largely below 1,500 bp. By contrast, the ARES (orange and blue) are constrained by the size of the GeneEE inserts. Notably, the IRs upstream of the top 10 % most highly expressed genes cluster at even shorter lengths than the genome-wide average.

**Figure 6.**
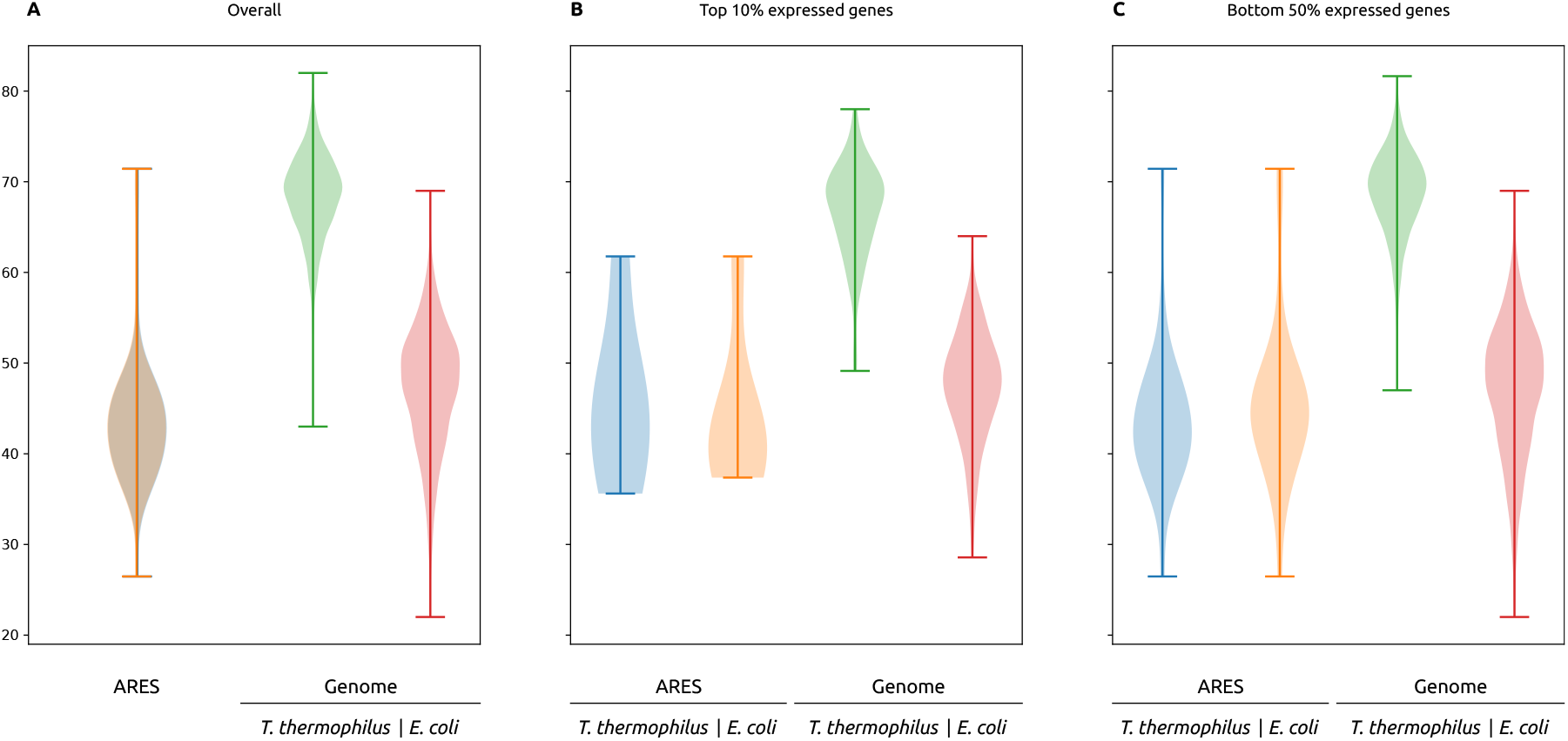
GC content distributions for *T. thermophilus, E. coli*, and ARES. Violin plots depict GC percentages for all ARES and genes at the corresponding genomes (A), the top 10% of expressed genes (B), and the bottom 50% of expressed genes (C).

**Figure 7.**
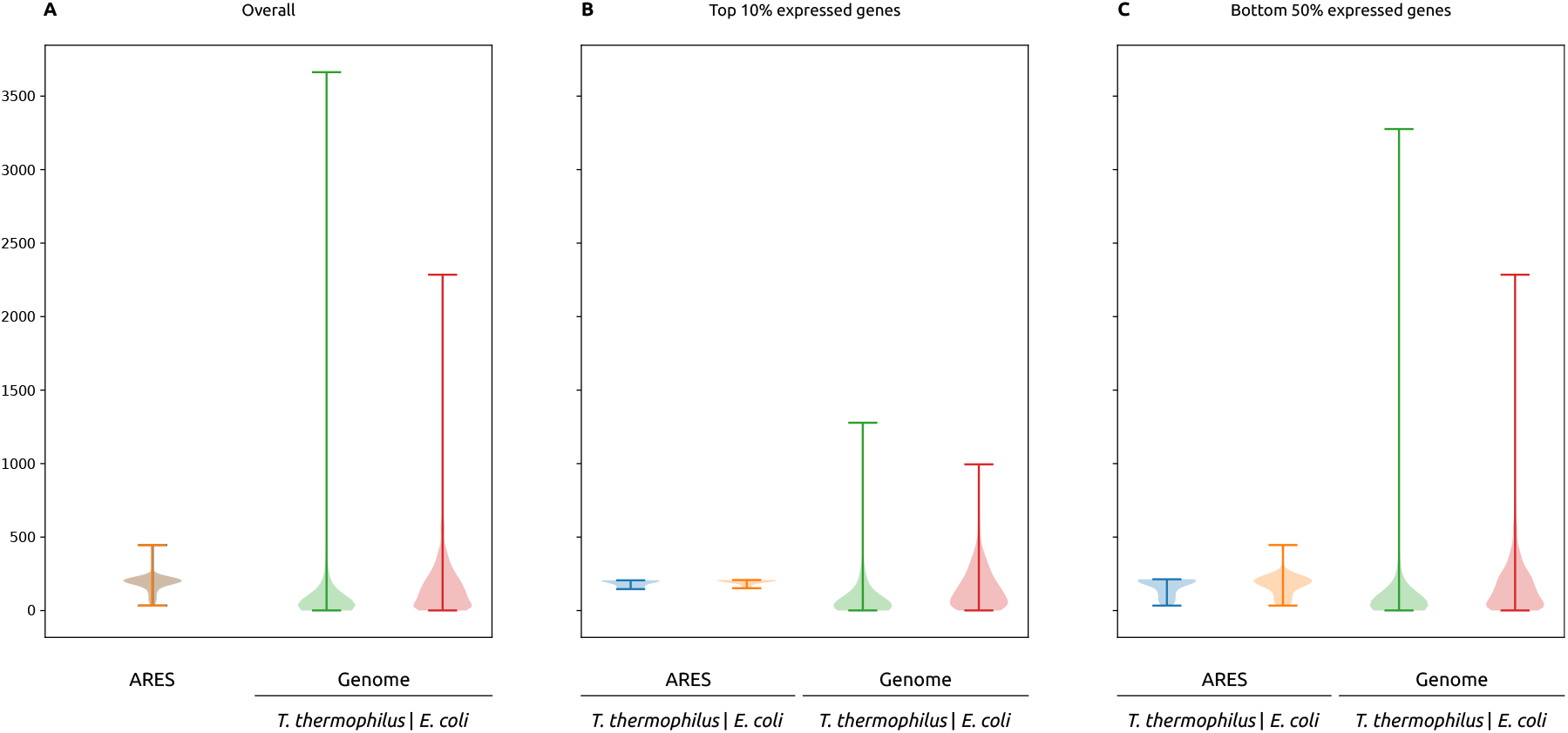
Intergenic region length distributions in *T. thermophilus, E. coli*, and their ARES. Violin plots compare intergenic region (IR) lengths for all ARES and genes the genome level (A), the top 10% of highly expressed genes (B), and the bottom 50% of weakly expressed genes (C). *T. thermophilus* (green) shows a broader range of IR lengths, some exceeding 3,000 bp, while *E. coli* (red) remains under 1,500 bp. The ARES (orange and blue) exhibit relatively short IR-like segments, reflecting their design for minimal essential promoter functionality.

Together, these results underscore the compact, high-GC nature of the *T. thermophilus* genome compared to that of *E. coli*. They also demonstrate that our synthetic ARESs diverge significantly from native promoters in both length and nucleotide composition (as confirmed by BLAST analyses). Such divergence is advantageous in synthetic biology, as it broadens the sequence space for uncovering novel, potentially stronger or more finely tunable transcriptional control elements.

## Discussion

In this study, we generated and functionally characterised 53 ARES for use in *T. thermophilus* and *E. coli*. Our results demonstrate that the GeneEE platform can rapidly generate a diverse set of regulatory elements without prior knowledge of native promoter architecture. The observed host-specific differences, where most ARES drive stronger expression in *T. thermophilus* compared to *E. coli*, highlight the importance of screening in the intended host system.

Notably, the ARES exhibit an average GC content of about 44%, markedly lower than that of the native promoters *P*_*slpA*_ and *P*_*nqo*_ (60–69 %). The BLAST analysis revealed no significant matches to known natural promoters, indicating that our artificial design accesses a seldom-explored region of sequence space in the thermophile. By avoiding homology to existing genomic elements, these non-natural sequences expand the available promoter repertoire for extremophilic hosts and pave the way for de novo regulatory engineering.

The functional characterisation of thermostable *β*-galactosidase and phytoene synthase further validates the utility of these ARES. The ability to modulate expression levels over a broad range is critical for optimising industrial processes. For example, the production of thermostable enzymes such as *β*-galactosidase is important in the dairy industry for the production of lactose-free products, while phytoene synthase is a key enzyme in carotenoid biosynthesis, a pathway of growing interest due to its applications in food colourants, nutraceuticals, and pharmaceuticals. Moreover, the potential to express heterologous pathways in thermophilic hosts could simplify downstream processing in high-temperature industrial processes.

In summary, our study enriches the synthetic biology toolkit for extremophiles by delivering a panel of novel, functionally validated ARESs, thereby laying the groundwork for both metabolic engineering strategies and the scalable production of high-value thermostable proteins in industrial settings.

## Author contributions statement

C.F.A.W. and R.L. conceived the study and involved in the design of experiments. C.F.A.W. and S.Z. carried out the experiments and wrote the initial draft. L.T. involved in plasmid DNA library construction experiments in *E. coli*. G.S.D. performed the bioinformatics analysis on the ARES and genomes of *E. coli* and *T. thermophilus*. All authors contributed to manuscript revision, read, and approved the submitted version.

## Acknowledgements

We thank Dr. José Berenguer (Universidad Autónoma de Madrid) who kindly provided the *T. thermophilus* strain used in this study. Open access was supported by the NTNU library.

